# DREADD activation of the lateral septum alters prosocial and antisocial behaviors, but not partner preferences in male prairie voles

**DOI:** 10.1101/2022.07.26.501564

**Authors:** Lindsay L. Sailer, Ashley H. Park, Abigail Galvez, Alexander G. Ophir

## Abstract

Although much has been written on the topic of social behavior, many terms referring to different aspects of social behavior have become inappropriately conflated and the specific mechanisms governing them remains unclear. It is therefore critical that we disentangle the pro- and anti-social elements associated with different forms of social behavior to fully understand the social brain. The lateral septum (LS) mediates social behaviors, emotional processes, and stress responses necessary for individuals to navigate day-to-day social interactions. The LS is particularly important in general and selective prosocial behavior (monogamy) but its role in how these two behavioral domains intersect is unclear. Here, we investigate the effects of chemogenetic-mediated LS activation on social responses in male prairie voles when they are 1) sex-naïve and generally affiliative and 2) after they become pair-bonded and display selective aggression. Amplifying neural activity in the LS augments same-sex social approach behaviors. Despite partner preference formation remaining unaltered, LS activation in pair-bonded males leads to reduced selective aggression while increasing social affiliative behaviors. These results suggest that LS activation alters behavior within certain social contexts, by increasing sex-naïve affiliative behaviors and reducing pair bonding-induced selective aggression with same-sex conspecifics, but not altering bonding with opposite-sex individuals.

## Introduction

For as long as people have studied social behavior, an underlying goal has been to understand the mechanisms that govern them. Neuroscientists with different backgrounds and expertise have attempted to define the core neural mechanisms that underlie social behavior, but defining a ‘social brain’ is as complicated as the behaviors that are presumably under its control. This is because social behavior takes so many different forms (affiliation, aggression, approach, consolation, mating, nurturing, play, etc.), each of which involves many different behavioral elements and interactions. Nevertheless, determining how and where the brain processes and shapes behavior in response to social factors is of great importance if we are to truly understand the nature and universality of social behavior.

O’Connell and Hoffmann (O’Connell & Hofmann, 2011) provided an expanded view of the neural control and modulation of social behavior, and introduced the idea of the social decision-making network (SDMN). The SDMN is comprised of interconnected neural structures that are heavily involved in the regulation of social behavior (O’Connell & Hofmann, 2011; A. G. Ophir, 2011). Contained within the SDMN are sub-circuits that are implicated in coordinating different aspects of social behavior, including social grouping (Goodson, Kelly, & Kingsbury, 2012; A. M. Kelly et al., 2011), aggression (Wong et al., 2016), parental care (Numan, 2020), and pair bonding (Johnson & Young, 2015; Prounis & Ophir, 2020). For example, work in prairie voles has defined a neural circuit that governs pair bonding, which has formed the basis for understanding the neurobiology of mammalian social affiliation and monogamy (reviewed in (Johnson & Young, 2015; Young & Wang, 2004)). Notably, the entirety of the pair bond neural circuit is encompassed by the SDMN (Prounis & Ophir, 2020). In contrast to affiliation and sociability, aggression and territoriality can be considered examples of the ‘dark side’ of social behavior. Nevertheless, aggression is a prevalent and fundamental aspect of social behavior that enables defense of resources, offspring, and mating partners. Not surprisingly, neural structures within the SDMN also play a prominent role in the regulation of aggression, and aggressive responses to intruders can be precisely modified through manipulations of sub-units of the SDMN (Wong et al., 2016). The lateral septum (LS) is a central node of the SDMN that plays a prominent role in the regulation of social behavior across vertebrates (O’Connell & Hofmann, 2011), and has been called the most unjustifiably ignored region of the human social brain (Lieberman, 2014). For instance, manipulations of the LS modify social grouping preferences in finches (Goodson et al., 2012; A. M. Kelly et al., 2011), aggression in mice (Wong et al., 2016), kin preferences in rats (Clemens, Wang, & Brecht, 2020), and pair bonding in prairie voles (Liu, Curtis, & Wang, 2001; Winslow, Hastings, Carter, Harbaugh, & Insel, 1993). Recently, Kelly *et al.* (A. M. Kelly, Ong, Witmer, & Ophir, 2020) demonstrated that social approach is associated with early-life social experience and epigenetic modification of the vasopressin receptor within the LS. Ultimately, the LS appears to impact pro-social and anti-social aspects of social behavior across contexts and species.

Prosocial and antisocial behaviors profoundly impact social monogamy, parental investment, and group structure (A. G. Ophir, 2011). Prairie voles (*Microtus ochrogaster*) form long-term pair bonds with mates, form intense attachments with offspring, display selective aggression to defend territories and mates, and exhibit bi-parental care for their young (Thomas & Birney, 1979; Winslow et al., 1993). Importantly, sexually inexperienced male and female prairie voles are rarely aggressive toward conspecifics and typically spend most of their time engaging in affiliative and investigative behaviors (Shapiro & Dewsbury, 1990). Mating in prairie voles facilitates the formation of a pair bond and creates a dramatic shift from engaging in general affiliation with familiar and novel conspecifics, to exhibiting selective affiliation with familiar conspecifics and aggression toward strangers. Specifically, pair bonded voles preferentially engage in partner- and offspring-directed affiliative behaviors, become profoundly territorial, and exhibit high levels of aggression toward unrelated males and females that trespass their territory. Thus, despite the common caricature that prairie voles are ‘highly social,’ they display a wide range of prosocial and antisocial behaviors according to their physiological and environmental states. Thus, the range of context-dependent prosocial and antisocial behavior in prairie voles raises the question: what is the neural basis for this flexibility in social behaviors that enables effective reproduction?

Capitalizing on resources and social interactions requires animals to appropriately adjust their behavioral output by calibrating the intensity and duration of prosocial and antisocial responses targeted at familiar or novel conspecifics (A. M. Kelly & Vitousek, 2017; A. M. Kelly & Wilson, 2020). The ability to achieve these adaptive behavioral responses to social stimuli and environmental constraints relies upon efficient transmission, appraisal, and processing of information via dynamic neural and molecular mechanisms (Prior, Bentz, & Ophir, 2022). Neuromodulation of the LS could serve this function. Sheehan and colleagues (T. P. Sheehan, Chambers, & Russell, 2004) have proposed that the LS appears to play a regulatory function for social behavior. They argue that it does so by integrating sensory stimuli and assessing their affective relevance and valence. The LS then conveys this information to other SDMN brain regions known to promote emotional states and/or for directing motivated behaviors. Thus, the LS can orchestrate behavioral responses by appropriately adjusting behavior to meet environmental demands (T. P. Sheehan et al., 2004).

Here, we hypothesize that the LS serves as a point of convergence for the contextual regulation of numerous aspects of social behavior central to reproduction. To this end, we assess the impact of LS chemogenetic stimulation on affiliation, aggression, and partner preference formation in male prairie voles. We evaluate the function of the LS before and after sexual experience with females to assess how LS activation impacts the shift from prosocial to selectively antisocial behavior that characterizes pair bonded prairie voles. As an often-overlooked component of the social brain (Lieberman, 2014; Prounis & Ophir, 2020), we aim to understand the modulatory role of the LS on regulating social behavior in male prairie voles as they transition from being generally sociable to socially selective. We conclude that the lateral septum is positioned to integrate social context-dependent information to produce flexible and appropriate behavioral responses.

## Results

To begin to evaluate the effects of LS behaviors, we injected an excitatory followed by intraperitoneal (i.p.) injections of 3mg/kg of compound C21 (hM3+C21, **Figure 1A**). Control adult males expressed the excitatory DREADD in the LS and were injected with saline (hM3+saline, **Figure 1A**). To control for off-target behavioral effects of C21 administration (Bonaventura et al., 2019), a second group of control adult males did not express the hM3 receptor in the LS and were injected with 3 mg/kg C21 (sham+C21, **Figure 1A**). Thus, we had three treatment groups, males with LS activation (hM3+C21) and two control groups (hM3+saline and sham+C21). Acute C21 and saline i.p. injections were administered 30 min before each behavioral test on days 16, 17, 20, and 21 (**Figure 1B**). To measure the effects of LS activation on sex naïve social approach and aggressive behaviors with age-matched and sex-matched stimulus males, subjects were assessed in the social approach test on day 16 and the resident intruder test on day 17, respectively. Then subjects cohabitated with sexually receptive females for 48 hours and the effects of LS activation on partner preference formation on day 20 were examined. After pair bonding, adult males were subjected to a second resident intruder test on day 21 to assess the effects of LS activation on pair bond-induced aggression. In summary, we found LS activation influenced both prosocial and selectively aggressive behaviors in adult males when they were pair bonded, but not when they were sex naïve. Notably, activation on adult male sex naïve and pair bonded social DREADD virus (AAV8-hSyn-hM3D-mCherry) into the LS, the socially selective partner preference was not affected by LS activation.

**Figure 1.**
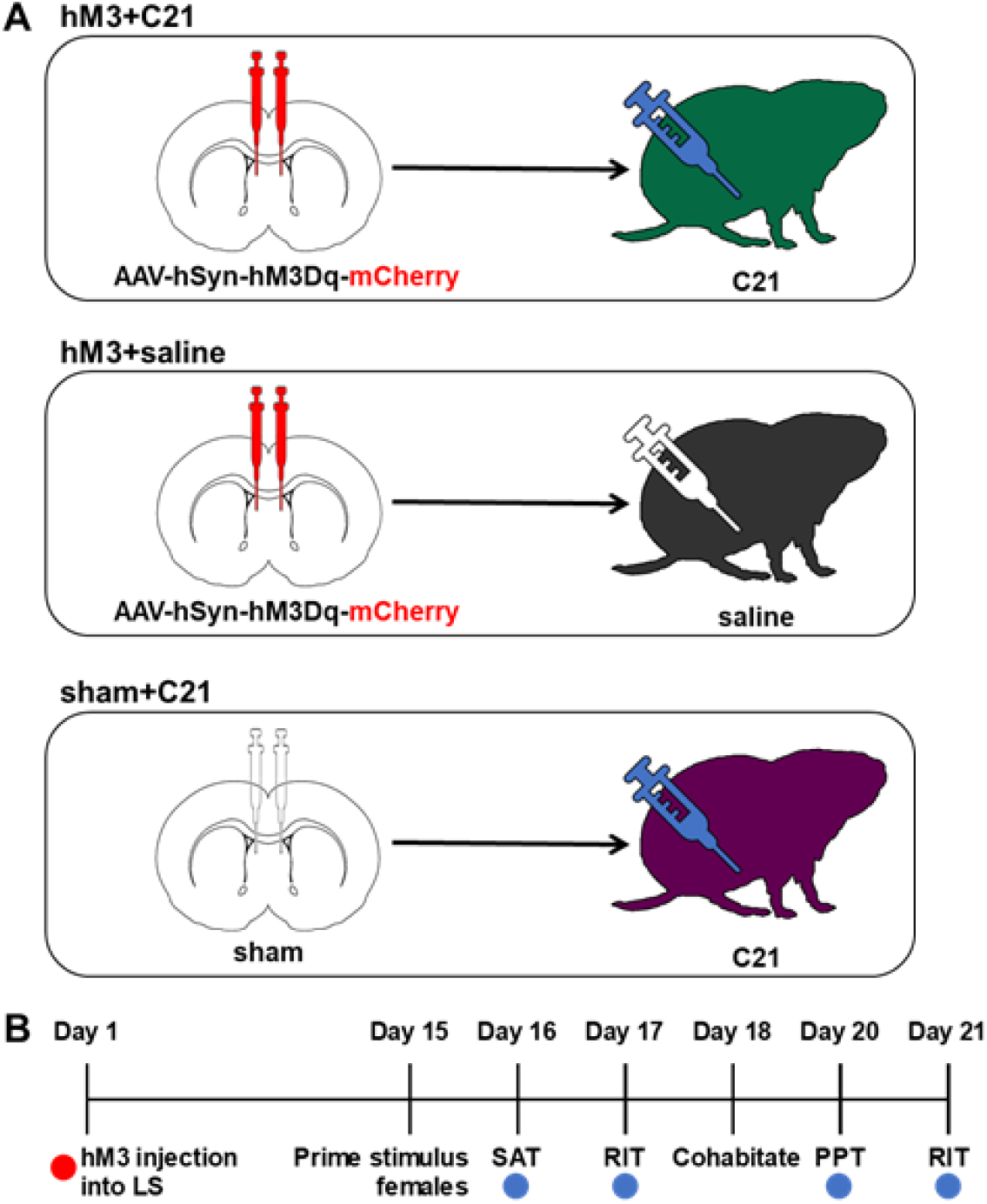
Enhancing neural activity in the lateral septum. (A) Schematic illustrating animal groups receiving AAV8-hSyn-hM3D-mCherry injections into the LS and 3 mg/kg C21 i.p. injections (hM3+C21, green); AAV8-hSyn-hM3D-mCherry injections into the LS and saline i.p. injections (hM3+saline, grey); or sham injections into the LS and 3 mg/kg C21 i.p. injections (sham+C21, purple). (B) Timeline of experimental design. LS, lateral septum; SAT, social approach test; RIT, resident intruder test; PPT, partner preference test. Red dot indicates when hM3Dq-mCherry or sham injection surgeries targeting the LS were performed. Blue dots indicate when 3 mg/kg C21 or saline (i.e., vehicle) were injected (i.p.) 30 min prior to each behavioral assessment.

### LS activation promotes sex naïve social approach

We first assessed behavioral responses to LS activation by evaluating social approach behaviors in sex naïve males on day 16 (**Figure 1B**). We found a significant interaction for the duration of time spent in the non-social and social zones of the testing apparatus (LMM, treatment x zone interaction: F_(2,64)_ = 5.35, *p* = 0.007; **Figure 2A**). LS activation significantly increased duration in the social zone relative to the non-social zone (*post hoc t*_35.3_ = −2.76, *p* = 0.009). LS activation (hM3+C21) did not alter the latency to social approach (treatment effect: F_(2,29)_ = 0.97, *p* = 0.39; **Figure 2B**) or frequency of zone transitions (treatment x zone interaction: F_(2,32)_ = 0.004, *p* = 0.10; **Figure 2C**). Distance moved (treatment effect: F_(2,29)_ = 3.19, *p* = 0.06; **Figure S1**) and velocity (treatment effect: F_(2,29)_ = 3.21, *p* = 0.06; **Figure S2**) were not significantly decreased by LS activation, although both demonstrated non-significant trends toward more movement among the hM3+saline control group, largely driven by three individuals (see supplementary material). These data indicate that activation of the lateral septum promotes social approach to a novel male conspecific, while locomotor behavior in sex naïve male prairie voles remained unaltered.

**Figure 2.**
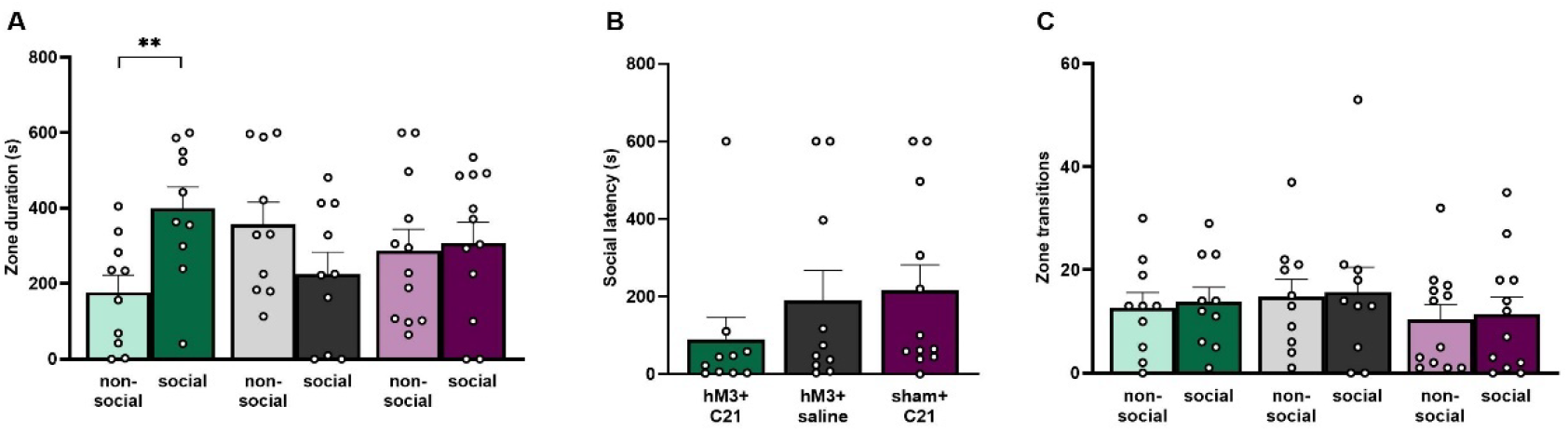
LS activation promotes sex naïve social approach. (A) Duration (s, seconds) of time spent in the non-social (light color bars) and social zones (dark color bars) of the social approach test apparatus by hM3+C21, hM3+saline, and sham+C21 subjects. (B) Latency to approach (s) stimulus animal by hM3+C21, hM3+saline, and sham+C21 subjects. (C) Frequency of zone transitions between non-social (light color bars) and social zones (dark color bars) by hM3+C21 (green bars), hM3+saline (grey bars), and sham+C21 (purple bars) subjects. Data are presented as mean ± SEM. Dots represent individual data. ** *p* < 0.01.

### LS activation does not alter partner preference formation

We investigated whether LS activation would alter the ability of males to form socially selective partner preferences. Across treatments, males spent more time in contact with the partner than the stranger (F_(1,58)_ = 20.98, *p* = 2.51×10^−5^, **Figure 3A**). Specifically, control males formed a significant partner preference with their mate (partner vs stranger: hM3+saline, *t*_32.4_ = 2.53, *p* = 0.008; sham+C21, *t*_32.4_ = 3.09, *p* = 0.002), as well as males whose LS was activated (partner vs stranger: hM3+C21, *t*_32.4_ = 1.93, *p* = 0.03). Moreover, partner preference indices (time with partner – time with stranger) for each treatment group (F_(2,26)_ = 0.18, *p* = 0.84; **Figure 3B**) and total time spent in contact with both stimulus animals (time with partner + time with stranger) for each treatment group (F_(2,26)_ = 1.04, *p* = 0.37; **Figure 3C**) were not significantly different, indicating that the preference for the partner was, the same for all three groups.

**Figure 3.**
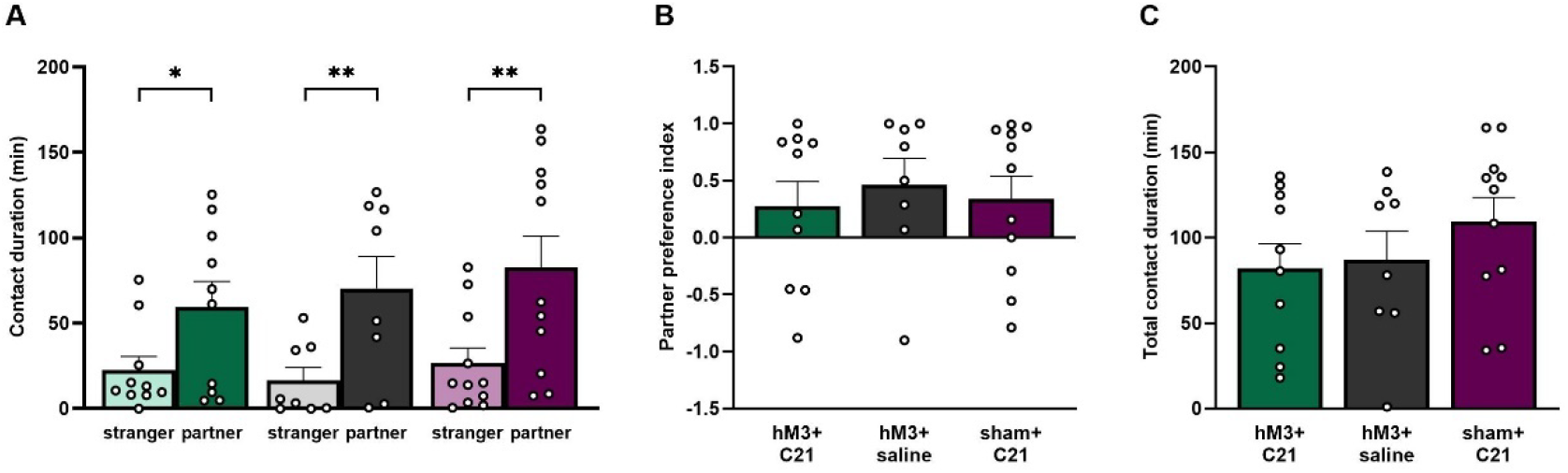
LS activation does not alter partner preference formation. (A) Contact duration (min, minutes) when hM3+C21 (green bars), hM3+saline (grey bars), and sham+C21 (purple bars) subjects spent time with the stranger (light color bars) or partner (dark color bars) stimulus females. (B) Partner preference index (contact duration with partner – contact duration with stranger) / (contact duration with partner + contact duration with stranger). (C) Total contact duration (min) with both the stranger and partner (contact duration with partner + contact duration with stranger). Data are presented as mean ± SEM. Dots represent individual data. * *p* < 0.05; ** *p* < 0.01.

Importantly, cage transitions were not affected by LS activation (treatment main effect: F_(2,26)_ = 0.16, *p* = 0.85; **Figure S3**). We interpret these data as additional evidence demonstrating that activation of the LS does not alter the ability for male prairie voles to form partner preferences. Furthermore, LS activation did not affect the overall preference for social engagement (**Figure 3C**), or locomotor behavior (**Figure S3**) in pair bonded males because total contact duration and cage transitions were similar between groups. Taken together, we conclude that LS activation did not alter overall opposite-sex social selectivity in male prairie voles.

### LS activation reduces pair bond-induced aggression

Males are disproportionately aggressive with strangers in the resident-intruder paradigm only after forming a pair bond, but not before (Winslow et al., 1993). We examined how LS activation affects aggressive, social, defensive, and non-social behaviors before (sex naïve; day 17) and after (day 20) pair bond formation in the resident-intruder test (**Figure 1B**), which we refer to as ‘bond status’.

Duration of attacks increased after forming a pair bond compared to when males were sex naïve (bond status, F_(1,32)_ = 7.59, *p* = 0.0001; **Figure 4A**). Importantly, *post hoc* analysis revealed that only control males (hM3+saline: *t*_35.3_ = 2.03, *p* = 0.05; and sham+C21: *t*_35.3_ = 2.97, *p* = 0.005) engaged in a greater duration of attack behaviors when they were pair bonded compared to when they were sex naïve (i.e., before they were bonded). In contrast, LS activation significantly inhibited pair bond-induced attack behaviors directed at the intruder when males were pair bonded compared to when they were sex naïve (hM3+C21: sex naïve vs pair bonded: *t*_35.3_ = −0.332, *p* = 0.74).

**Figure 4.**
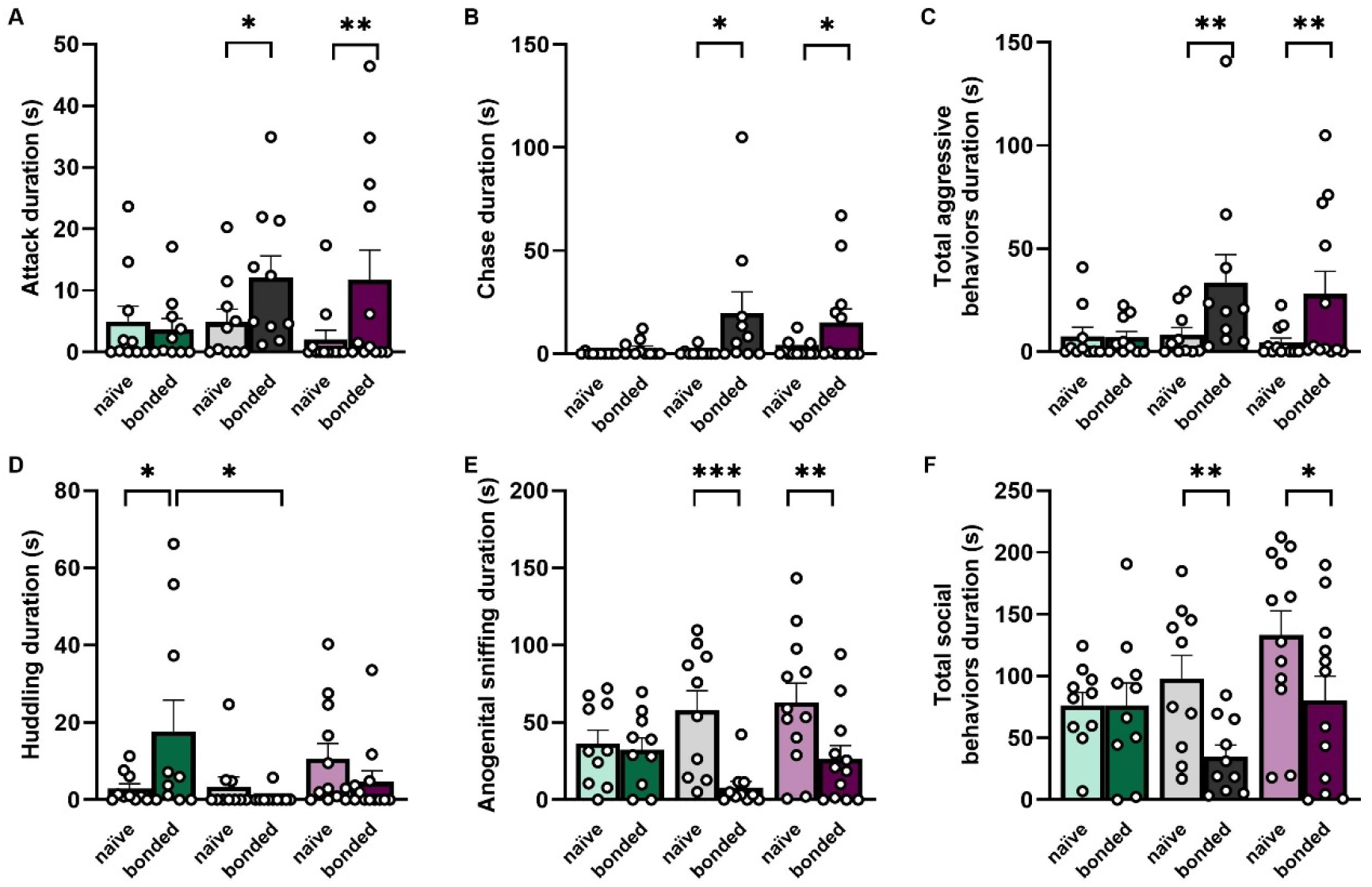
LS activation reduces pair bond-induced aggression and promotes prosocial behaviors. Duration (s, seconds) of attacks (A), chasing (B), total aggressive behaviors (C, sum of the duration of attack, chase, and pounce behaviors), huddling (D), anogenital sniffing (E), and total social behaviors (F, sum of the duration of huddling, anogenital sniffing, flank sniffing, and nose-to-nose sniffing behaviors) directed at the intruder by the resident hM3+C21 (green bars), hM3+saline (grey bars), and sham+C21 (purple bars) subjects when they are sex naïve (light color bars) and pair bonded (dark color bars) during the resident intruder test. Data are presented as mean ± SEM. Dots represent individual data. * *p* < 0.05; ** *p* < 0.01.

The latency to attack the intruder was affected by LS activation (treatment, F_(2,32)_ =5.75, *p* = 0.007) and bond status (F_(1,32)_ = 19.03, *p* = 0.0001). The interaction effect fell short of our significance threshold (treatment x bond status, F_(2,32)_ = 2.92, *p* = 0.07). *Post hoc* analysis revealed that, as expected, sex naïve males did not differ in latency to attack (hM3+C21 vs hM3+saline: *t*_69.3_ = −0.97, *p* = 0.60; hM3+C21 vs sham+C21: *t*_69.3_ = 1.88, *p* = 0.15; hM3+saline vs sham+C21: *t*_69.3_ = 0.87, *p* = 0.66). However, once pair bonded, hM3+saline males significantly decreased their latencies to attack in comparison to when they were sex naïve (*t*_35.3_ = −4.18, *p* = 0.0002), showing the expected pattern of post-bond selective aggression (Thomas & Birney, 1979; Winslow et al., 1993). The sham+C21 males also tended to decrease attack latency, but this decrease was not significant (*t*_35.3_ = −1.58, *p* = 0.12). In contrast, LS activation inhibited post-bond selective aggression; the latency to attack the intruder remained long in hM3+C21 males when they were sex naïve and pair bonded (*post hoc t*_35.3_ = −1.36, *p* = 0.18). Moreover, LS-activated males (hM3+C21) were slower to attack the intruder than hM3+saline control males once they were pair bonded (*t*_69.3_ = −3.59, *p* = 0.002), and tended (non-significantly) to be slower to attack than pair bonded sham+C21 males (*t*_69.3_ = 1.96, p = 0.13).

Subject males increased the instances of aggressive chases overall as they transitioned from sex naïve to pair bonded (F_(1,32)_ = 9.41, *p* = 0.004; **Figure 4B**). Specifically, males from both control groups chased intruders for more time when they were pair bonded compared to when they were sex naïve (hM3+saline: *t*_35.3_ = 2.71, *p* = 0.01; sham+C21: *t*_35.3_ = 2.05, *p* = 0.05). However, LS activation significantly inhibited pair bond-induced intruder chasing behaviors. That is, hM3+C21 males did not differ in time spent chasing intruders when they were pair bonded compared to when they were sex naïve (*t*_35.3_ = 0.34, *p* = 0.74). Duration of pouncing did not yield any significant main effects (treatment, F_(2,64)_ = 1.07, *p* = 0.35; bond status, F_(1,64)_ = 0.40, *p* = 0.53) or a significant interaction of treatment x bond status (F_(2,64)_ = 0.69, *p* = 0.50).

Finally, we examined how LS activation impacted the proportion of time subjects spent engaging in total aggressive behaviors (sum of the duration of attack, chase, and pounce behaviors). Linear mixed-model analysis revealed a significant main effect of bond status (F_(1,32)_ = 10.63, *p* = 0.003; **Figure 4C**). *Post hoc* analysis revealed that control males aggressively targeted the intruder significantly more when they were pair bonded compared to when they were sex naïve (hM3+saline: *t*_35.3_ = 2.75, *p* = 0.009; sham+C21: *t*_35.3_ = 2.78, *p* = 0.009). Unlike the two control groups, LS activation inhibited the normal expression of pair bond-induced aggressive behaviors in hM3+C21 males (*t*_35.3_ = −0.06, *p* = 0.95). Taken together, these results indicate that LS activation inhibited selective aggression in pair bonded males.

### LS activation increases prosocial behavior among pair bonded males

The reduction of aggressive behaviors in LS-activated males was accompanied by an increase in social behaviors. Specifically, we found a significant interaction between treatment and bond status (F_(2,32)_ = 4.46, *p* = 0.02; **Figure 4D**) when examining the duration of huddling behaviors during the resident intruder test. Notably, *post hoc* analyses revealed that LS activation significantly increased the duration of huddling with an intruder when hM3+C21 males were pair bonded compared to when they were sex naïve (*t*_35.3_ = 2.60, *p* = 0.01). In contrast, control males did not differ in their duration of huddling when they were sex naïve or pair bonded (hM3+saline: *t*_35.3_ = −0.51, *p* = 0.61; sham+C21: *t*_35.3_ = −1.14, *p* = 0.26). Moreover, the duration of huddling in pair bonded LS-activated (hM3+C21) males was significantly longer than pair bonded hM3+saline control males (*t*_70.6_ = −2.98, *p* = 0.01), and pair bonded sham+C21 control males (*t*_70.6_ = 2.40, *p* = 0.05).

Anogenital sniffing, a common marker of prosocial investigation, during the resident intruder test also showed a significant main effect of bond status (F_(1,32)_ = 21.81, *p* = 5.17×10^−5^; **Figure 4E**) and a significant interaction between treatment and bond status (F_(2,32)_ = 4.19, *p* = 0.02). *Post hoc*analysis revealed that LS activation maintained the high instances of sex naïve anogenital sniffing after the bond was formed (hM3+C21: *t*_35.3_ = −0.34, *p* = 0.73). Control males, however, demonstrated the anticipated reduction in prosocial anogenital sniffing with the intruder once they had established a bond (hM3+saline: *t*_35.3_ = −4.12, *p* = 0.0002; sham+C21: *t*_35.3_ = −3.31, *p* = 0.002).

Lastly, we examined how LS activation impacted the total time subjects spent engaging in social behaviors (sum of the duration of huddling, anogenital sniffing, flank sniffing, and nose-to-nose sniffing behaviors) when males were sex naïve and pair bonded. Our results demonstrated a significant main effect of bond status (F_(1,32)_ = 9.77, *p* = 0.004; **Figure 4F**) on total social behaviors. *Post hoc* analysis revealed that control males, as expected, spent less time engaging in social behaviors with an intruder when they were pair bonded compared to when they were sex naïve (hM3+saline: *t*_35.3_ = −2.73, *p* = 0.0001; sham+C21: *t*_35.3_ = −2.51, *p* = 0.02). In contrast, LS activation maintained the expression of prosocial behaviors with a stranger after bonding (hM3+C21: *t*_35.3_ = 0.006, *p* = 0.99). Taken together, results from comparisons of aggressive and prosocial behaviors in the resident-intruder paradigm indicate that LS activation interferes with the selective aggression typical of bonded male prairie voles, and actually promotes prosocial behaviors with intruders.

## Discussion

Sex naïve prairie voles seldomly exhibit aggressive behaviors and readily engage socially with novel conspecifics, but they become highly socially selective after mating (Shapiro & Dewsbury, 1990). Here, we demonstrate that LS activation eliminated agonistic elements of this social selectivity and enhanced sex naïve-like preferences for social novelty in pair bonded males. Remarkably, LS-activated males continued to demonstrate a partner preference, indicating that social selectivity for a partner remained intact and was overlaid on top of the enhancement of general prosocial behaviors. Non-social locomotor behaviors were unaffected by chemogenetic stimulation of the LS, indicating that enhanced LS activation specifically modulated prosocial and antisocial behaviors. Taken together, we demonstrate that the LS effectively regulates prosocial and antisocial behaviors in a context-dependent manner consistent with life-history transitions in reproductive behaviors.

The transition from being a single (un-bonded) male to a pair bonded socially monogamous male is accompanied by a suite of characteristic changes in behavior. Single males (referred to as ‘wanderers’ in nature) are non-territorial and typically inhabit expansive home ranges that intrude into the home ranges of many other conspecifics (Madrid, Parker, & Ophir, 2020). The high rates of home range overlap appear to be indicative of indiscriminate prosocial attraction to others. Once bonded, the behavioral demands on male prairie voles change. Bonded males (referred to as ‘residents’ in nature) are territorial, appear to mate guard, and become (at least initially) selectively aggressive to conspecifics other than their partner and offspring. The reproductive success of wanderers is less than, or possibly equal to, that of their pair bonded resident counterparts (A. G. Ophir, Phelps, S. M., Sorin, A. B., Wolff J. O., 2008; Shuster, Willen, Keane, & Solomon, 2019) suggesting that males benefit reproductively to a greater extent when bonded. Notably, the putative reproductive advantage for residents is directly associated with this behavioral shift, in which males go from being indiscriminately social and relatively non-aggressive, to being selectively social with partners and selectively aggressive to other adults. Our study provides insight into the mechanistic control of this specific behavioral shift, implicating LS as a major node in the network that modulates these forms of social behavior. Specifically, we believe our data support the hypothesis that the LS functions to shift the balance between general affiliation and social selectivity in a context-dependent manner (Luo, Tahsili-Fahadan, Wise, Lupica, & Aston-Jones, 2011; Sartor & Aston-Jones, 2012; T. P. Sheehan et al., 2004). In this case, such a shift enables the behavioral transition associated with being single to being bonded.

The neurobiology of pair bonding has benefited greatly from studies using prairie voles to uncover the specific mechanisms that modulate this important process. Pair bonding is rare among mammals (Kleiman, 1977), but mechanisms that control prairie vole bonding (including the actions of vasopressin and oxytocin within the LS (Liu et al., 2001)) seem to parallel the mechanisms that govern human social attachment (Insel & Young, 2001; Young & Wang, 2004). The LS has long been appreciated as a central node within pair bonding neural circuitry (Young & Wang, 2004). Yet the role of the LS in this process has not been terribly clear. One of the first pieces of evidence linking LS neural activation with reproductive history and mating-induced aggression in male prairie voles was demonstrated by Wang and colleagues (Wang, Hulihan, & Insel, 1997), where males exposed to females showed elevated Fos induction in the LS in comparison to males that never interacted with a female, irrespective of mating. Additionally, males that had prior experience with a female exhibited higher Fos staining in the LS in response to a male intruder relative to males that had no prior experience with a female (Wang et al., 1997). At the time, it was difficult to determine the physiological significance of Fos induction in the LS (i.e., activation vs inhibition). Evidence from our chemogenetic manipulations likely indicates that the LS was inhibited in the males that showed mating-induced aggression toward male intruders and Fos induction in the LS.

One hypothesis aimed at explaining the functional role of the LS in bonding is that it facilitates social recognition (Bielsky, Hu, Ren, Terwilliger, & Young, 2005), which can be paired with highly valanced social reward during mating, thereby enabling animals to associate selective preferences for a particular partner over all others (Walum & Young, 2018). Indeed, the LS (and oxytocin and vasopressin acting therein) have been frequently associated with differences in social recognition and discrimination, supporting this interpretation. Furthermore, like the other behaviors just discussed, prairie vole social recognition is altered after a bond has formed. Zheng *et al.* (Zheng, Foley, Rehman, & Ophir, 2013) showed that sex naïve adult male prairie voles fail to distinguish female conspecifics but effectively discriminate between males. This indiscriminate social recognition of females but clear ability to discriminate between males occurs when males are single. In nature, single males are described as wanderers - a pivotal period of life when it is necessary to be indiscriminate of mate choice (i.e., find any willing partner) while taking note of and being equipped to avoid potentially aggressive resident males and wanderer competitors. Once males establish a bond and form territories, a shift in cognition occurs where they now discriminate between females (Blocker & Ophir, 2015). The cognitive shift for social discrimination that is associated with pre- and post-bonding reproductive status presumably enables males to distinguish among conspecifics and act appropriately in prosocial (selective affiliation) or antisocial (selective aggression) interactions. Taken together, the LS is well positioned to enable numerous shifts in cognition and behavior that facilitate this key change in life-history stages.

The shift between pro-social and antisocial behaviors within prairie voles that we have reported is consistent with previous work that has focused on Estrildid finches, an all-monogamous genus of birds. Kelly, Goodson and colleagues (A. M. Kelly et al., 2011) elegantly demonstrated that the medial bed nucleus of the stria terminalis (BSTm) and the LS modulate social grouping based on the impact of vasotocin (VT, the non-mammalian form of vasopressin). In fact, the neural expression and function of VT directly relates to sociality in solitary and gregarious species of Estrildid finches (Goodson et al., 2012; A. M. Kelly et al., 2011). Manipulation of the BSTm-LS circuit in Estrildid finches alters social grouping behaviors, suggesting that VT impacts preferences for social grouping independently of mating system (Goodson et al., 2012). These results in birds strongly parallel our results in voles, in which DREADD-mediated LS activation eliminated selective aggression and promoted pro-social behaviors in both a social approach test and a resident-intruder paradigm, while leaving the pair bond intact. This important caveat highlights the notions that 1) the LS impacts some general aspects of pro- and antisocial behavior independently of other specific forms of social affiliation (i.e., bonding and mate preferences), 2) that it has species-specific functions that are generally similar but also particular to the life-history of different species (e.g., Estrildid finches do not undergo the life-history shifts in sociability to which prairie voles are subjected), or 3) both.

Yet, to assume that the LS is so specialized to control only social grouping behaviors or social recognition, for example, ignores the evidence that has implicated it in numerous forms of social behaviors and closely associated cognitive behaviors, such as kin preference (Clemens et al., 2020), social attachment to mates (Liu et al., 2001; Wang, 1995), social memory (Dantzer, Koob, Bluthe, & Le Moal, 1988; Everts & Koolhaas, 1999), social fear during lactation (Menon et al., 2018), juvenile play (Veenema, Bredewold, & De Vries, 2013), social approach (A. M. A. M. Kelly et al., 2020), and aggression (Leroy et al., 2018; Veenema, Beiderbeck, Lukas, & Neumann, 2010; Wong et al., 2016). These studies have beautifully demonstrated the modulatory role the LS plays in many discrete aspects of social behavior. For example, lesioning the LS and LS-GABA_A_ receptor activation causes irritability and aggression in mice (Slotnick, McMullen, & Fleischer, 1973; Wong et al., 2016) and hamsters (McDonald, Markham, Norvelle, Albers, & Huhman, 2012). Further, recent work by Clemmens *et al.* (Clemens et al., 2020) has shown that lesioning the LS disrupts age-dependent kin preferences in young rat pups. Instead of lesioning or inactivating the LS, we took a complimentary approach to chemogenetically activate the LS and examine its effect on prosocial behaviors and aggression in sex naïve male prairie voles, and later after they had become pair bonded. Notably, suppression of pair bond-induced aggression and promotion of social behaviors via LS activation in male prairie voles is congruous with previous work using optogenetics to activate LS neurons in sexually experienced male mice during resident intruder testing (Wong et al., 2016). Wong and colleagues (Wong et al., 2016) demonstrated that optogenetic stimulation of the LS in male mice significantly decreases the latency to attack, decreases the duration of attack behaviors, and increases the amount of time spent investigating a male intruder. Moreover, when the intruder is a female, LS optogenetic activation suppressed aggression, decreased the latency to stop mounting, and reduced the duration of time spent mounting (Wong et al., 2016).

A broader view of the function of the LS is that it plays a key role in modulating context-specific motivational states (Luo et al., 2011; Sartor & Aston-Jones, 2012; T. P. Sheehan et al., 2004). Indeed, the LS might function to modulate behavioral responses appropriate to particular environmental stimuli or internal life-history states through its connections within a larger network of brain areas, due to its cellular heterogeneity, or both (Rizzi-Wise & Wang, 2021). Several excellent candidate circuits have been characterized, most of which are directly or indirectly associated with understanding the general control of social behavior (A. G. Ophir, 2017; Prounis & Ophir, 2020; T. Sheehan, Numan, M., 2000; T. P. Sheehan et al., 2004). We believe that selective DREADD-mediated activation of the LS in our study tapped into and disrupted the dynamics within such circuitry and cellular dynamics that normally function to enable prairie voles to adjust behavior appropriately to account for important life-history shifts necessary to maximize their reproductive success. In this way, we have potentially provided evidence supporting the notion that the LS modulates context-specific motivational states in pro- and antisocial behavior that are dictated by bonding status.

In sum, we leveraged the social bonding nature of prairie voles to examine the impact that the lateral septum has on altering both prosocial and antisocial behaviors during a life-history transition from being single to being paired within the same individuals. Our study suggests that DREADD-mediated activation of the LS promotes prosocial behaviors and inhibits pair bonding-induced selective aggression, but does not affect the critical ability to form selective bonds and partner preferences. The LS may act as a nexus of prosocial and antisocial behaviors in male prairie voles, permitting physiological states and environmental demands to influence social phenotypes and reproductive decision-making. We previously argued that mating systems could be viewed as an independent ‘behavioral axis’ that is orthogonal to sociability (A. G. Ophir, 2011). Notably, the LS is critical for the expression of both monogamous pair bonding and sociability/aggression, suggesting that it could serve as a point of communication between the sub-networks that govern different forms of social behavior (A. G. Ophir, 2017; T. Sheehan, Numan, M., 2000; T. P. Sheehan et al., 2004) to facilitate the greater function of the social brain (Prounis & Ophir, 2020). In prairie voles, the shift from general sociability to social selectivity is a natural consequence of mating, which serves to guard mates and is an essential element of pair bonding. To our knowledge this is the first study to examine how chemogenetic control of the LS can shift the balance between general affiliation and social selectivity in male prairie voles as they transition from being sexually inexperienced and generally affiliative, to forming a pair bond by displaying social selectivity and territorial aggression. Our study advances understanding of the potential role that the LS takes on altering both prosocial and antisocial behaviors during a life history transition in reproductive state within the same individuals.

## Materials and Methods

### Animals

Male and female prairie voles used in this study were produced in our breeding colony at Cornell University, from breeding pairs that were offspring of wild caught animals captured in Champagne County, Illinois, USA. All subjects were unrelated, and sexually mature virgin males between 90 and 120 days old. Animals were weaned and housed with littermates on postnatal day (PND) 21, and then housed with same-sex littermates after PND 42-45. All animals received rodent chow (Laboratory Rodent Diet 5001, LabDiet, St. Louis, MO, USA) and water *ad libitum,* and were maintained under standard laboratory conditions (14L:10D cycle, lights on at 08:00, 20 ± 2 °C) in transparent polycarbonate cages (46.5 × 25 × 15.5 cm) lined with Sani-chip bedding and provided nesting material. All experimental procedures were conducted and approved by the Institutional Animal Care and Use Committee (IACUC) of Cornell University (2013-0102) and were in accordance with the guidelines set forth by the National Institutes of Health.

### Viral vector and stereotaxic surgery

The AAV8-hSyn-hM3D(Gq)-mCherry excitatory DREADD (hM3 for short), a gift from Bryan Roth (Addgene plasmid # 50474), was diluted to 1 × 10^12 vg/mL in sterile 0.1M PBS and stored in 5 μl aliquots at −80°C until the day of use. Before surgery, male subjects were anesthetized with 1.5-2% isoflurane mixed with pure oxygen (1 L//min) and fixed in a stereotaxic apparatus (Kopf Instruments). The scalp area was scrubbed with povidone-iodine (Purdue Products), and ophthalmic ointment (Henry Schein) was applied to the eyes. Subjects received either bilateral injections of the excitatory DREADD or sterile 0.1M PBS into the LS (+ 0.85 mm anterior, ± 0.55 mm lateral, and − 3.80 mm ventral relative to bregma; with bregma and lambda deviating up to ± 0.15 mm on the D/V axis). The virus suspended in PBS or PBS alone, was delivered at a volume of 300 nL/side using a 1.0 μL syringe (Hamilton Laboratory Products, Reno, NY), at a rate of 75 nL/min (**Figure 1A**). Following surgery, subjects were returned to their cages and administered acetaminophen orally (300 mg/kg body weight) in drinking water for 72 h and allowed to recover for an additional 13 d before behavioral experiments.

### Behavioral procedures

Sixteen days after administering DREADDs to the LS (day 0), all subjects underwent a social approach test to measure sex naïve same-sex affiliation. Four days later (20d post-surgery), animals were subjected to a partner preference test to measure their ability to form pair bonds with opposite-sex conspecifics. Two resident intruder tests were administered to subjects to measure sex naïve aggression (on day 17, before a bond could be established) and pair bond-induced aggression (on day 21, after a bond could be established). **Figure 1B** presents a schematic timeline for surgical and behavioral procedures. To activate the LS, subjects expressing the hM3 receptor were injected with 3 mg/kg of the DREADD agonist compound 21 (C21: Hello Bio, HB6124); control animals received saline vehicle. Thirty minutes before each behavioral test began (the social approach, resident intruder, and partner preference tests), subjects were treated with an acute injection of either 3 mg/kg C21 (hM3+C21) or saline (hM3+saline). A third group of subjects received 0.1M PBS injections into the LS to control for the injection of the hM3 DREADD viral vector and were acutely injected with 3 mg/kg C21 (sham+C21) 30 min before each behavioral test. This control group enabled assessment of potential C21-induced off-target effects in the absence of the hM3 receptor (Bonaventura et al., 2019). C21 was dissolved in sterile saline and injected intraperitoneally (i.p.) through a 26-gauge needle and 1 mL syringe. All animals were included in the analyses (n = 10 animals/treatment group).

#### Social approach test

Animals were introduced and allowed to acclimate to the social approach test (SAT) apparatus (20 × 40 × 28 cm) for 30 min. After acclimation, a doorway separating the testing chamber from a stimulus presentation box (10.06 cm^3^) containing an unfamiliar, age-matched, same-sex conspecific was unblocked, exposing the subject to the stimulus male conspecific. The stimulus chamber was separated from the testing chamber with a wall containing 13 holes (1.27 cm diameter) allowing for visual, auditory, and olfactory contact between the two animals. Tests were video recorded for a 10 min trial and the duration in the social zone and non-social zone, latency to approach the stimulus chamber, zone transitions, distance moved, and velocity were quantified. We quantified the latency to social approach as the difference in time from the start of the test until the nose of the subject was within 3 cm of the stimulus chamber.

#### Cohabitation and partner preference test

All subjects were pair-housed with sexually primed females for 48 hours before the partner preference test (PPT) was performed (**Fig 1B**). Animals were not treated with i.p. injections of C21 or saline during this cohabitation period. To induce sexual receptivity, all stimulus females were exposed to dirty bedding from non-sibling male cages for three consecutive days prior to pairing with a subject male (Carter, Getz, Gavish, McDermott, & Arnold, 1980). The PPT is a classic choice test paradigm in which a subject is placed in the center of a three-chambered apparatus (106.7 × 50.8 × 30.5cm), comprised of a central ‘neutral chamber’ (45.7 × 50.8 × 30.5 cm) and two ‘side chambers’ (27.9 × 50.8 × 30.5 cm). Off-set doorways allow the stimulus animal unrestricted access to the entire apparatus. The female used for cohabitation (i.e., “partner”) is secured to one side chamber and a novel female (i.e., “stranger”) is secured to the other side chamber using lightweight chains attached to zip-tie neck collars that are attached to the wall of each side chamber. Stimulus animals adapt quickly to the tethers and collars, and can engage in the full range of natural behaviors. Male subjects are allowed to freely explore the apparatus and interact with either of the two females for 3 hours while being video recorded. A trained experimenter blind to the treatment groups quantified the contact duration between the subjects and both their partners and strangers. Contact duration was calculated as time spent huddling, sniffing, and grooming the partner or stranger. This measure was used to calculate the total contact duration and a partner preference index (contact duration with partner – contact duration with stranger/ total contact duration with partner and stranger). A partner preference is defined as when the subject spends more time in side-by-side contact with the partner compared to the stranger (Young & Wang, 2004).

#### Resident intruder test

Levels of selective aggression exhibited by sex naïve and pair bonded males were examined using the resident intruder test (RIT) on testing days 17 and 21, respectively (**Fig 1B**). On day 17, the littermate was removed from the home cage and the resident’s interactions with an age-matched sex naïve male intruder was observed for 5 min. On day 21, the female partner was removed from the home cage and interactions between the resident male with a novel age-matched sex naïve male intruder were observed for 5 min. Duration of aggressive behaviors (attack, chase, and pounce) and prosocial behaviors (anogenital sniffing, nose-to-nose sniffing, flank sniffing, and huddling) were scored by a trained experimenter blind to the treatment groups.

### Confirmation of viral expression

Voles were transcardially perfused with cold 0.1M PBS, followed by 4% paraformaldehyde (PFA) in PBS under deep anesthesia. Brains were extracted and post-fixed for 24 h in 4% PFA, cryoprotected in 30% sucrose for 48 h, then frozen and stored in cryoprotectant at −80C°. Coronal sections (40 μm) containing the LS were cryosectioned (Leica Cryostat CM 1950) and collected for confirmation of DREADD expression via visualization of the mCherry fluorescent tag under a 10x objective (Leica DM550 B).

### Quantification and statistical analysis

All behavioral procedures were manually scored by an observer blind to treatment using Noldus Ethovision XT 13 (Noldus, Leesburg, VA, USA), Noldus Observer XT 11, or BORIS 7.9.7. Data were analyzed with RStudio (version 1.2.1335) using a linear mixed-model (LMM) framework with the packages lme4 (Bates, 2022), emmeans (Lenth, 2022), and lmerTest (Kuznetsova, 2020). Significant interactions or significant main effects (α ≤ 0.05) were followed by two-tailed Tukey’s *post hoc* test. For the partner preference test, a one-tailed (right sided) Tukey’s *post hoc* test was used because of our *a priori* assumption that subjects would exhibit a partner preference. For comparisons between treatment, data were tested for normality (Shapiro-Wilk) and equal variance. If data were not normally distributed, a Wilcoxon rank sum test was performed. Figures were created using Prism version 9.0.1.151 (GraphPad Software, San Diego California USA) and all data are presented as the means ± SEM.

Behaviors from the social approach test were analyzed to compare the effects of treatment between subjects (hM3+C21 vs hM3+saline vs sham+C21) on zone duration (non-social vs social), latency to social approach, zone transitions, distance moved, and velocity. Behaviors from the partner preference test were analyzed to compare the effects of treatment between subjects (hM3+C21 vs hM3+saline vs sham+C21) on contact duration (partner vs stranger), partner preference index, total contact duration, and cage transitions. Behaviors from the resident-intruder tests were analyzed to compare the effects of treatment between subjects (hM3+C21 vs hM3+saline vs sham+C21) and bonding status within subjects (sex naïve vs pair bonded) on social and aggressive behaviors.

## Acknowledgments

We thank Ophir lab members for their helpful feedback on the experiments. We would like to acknowledge the assistance of James Booth from the Cornell Statistical Consulting Unit during data analysis.

## Author Contributions

Conceptualization: LLS, AGO. Methodology: LLS, AGO. Investigation: LLS, AHP, AG. Visualization: LLS. Supervision: LLS, AGO. Writing—original draft: LLS, AGO. Writing—review & editing: LLS, AHP, AG, AGO.

## Funding

National Institutes of Health grant R01HD079573 (AGO)

Cornell University College of Arts and Sciences research support (AGO)

## Supplementary Material

**Figure S1.**
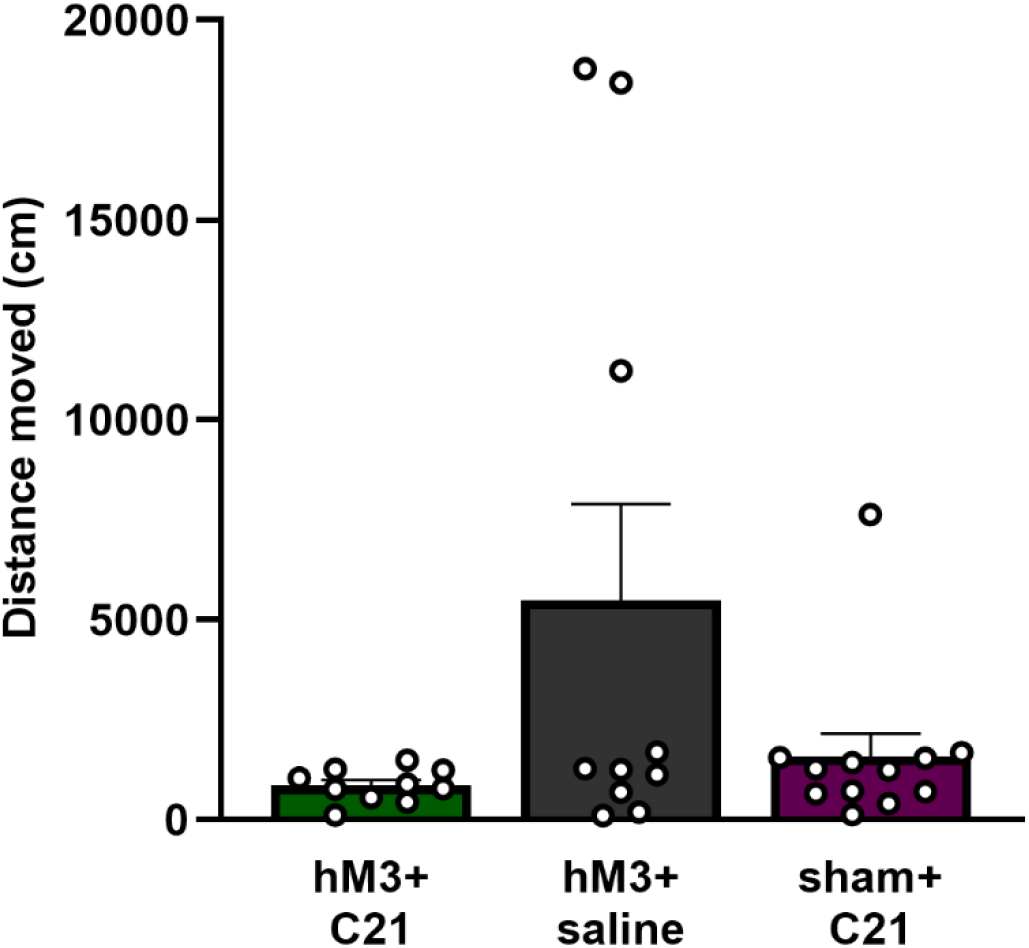
LS activation does not affect distance moved. Distance moved (centimeters; cm) by the hM3+C21 (green bars), hM3+saline (grey bars), and sham+C21 (purple bars) subjects during the social approach test. Data are presented as mean ± SEM. Dots represent individual data. (LMM, treatment effect: F_(2,29)_ = 3.19, p = 0.06).

**Figure S2.**
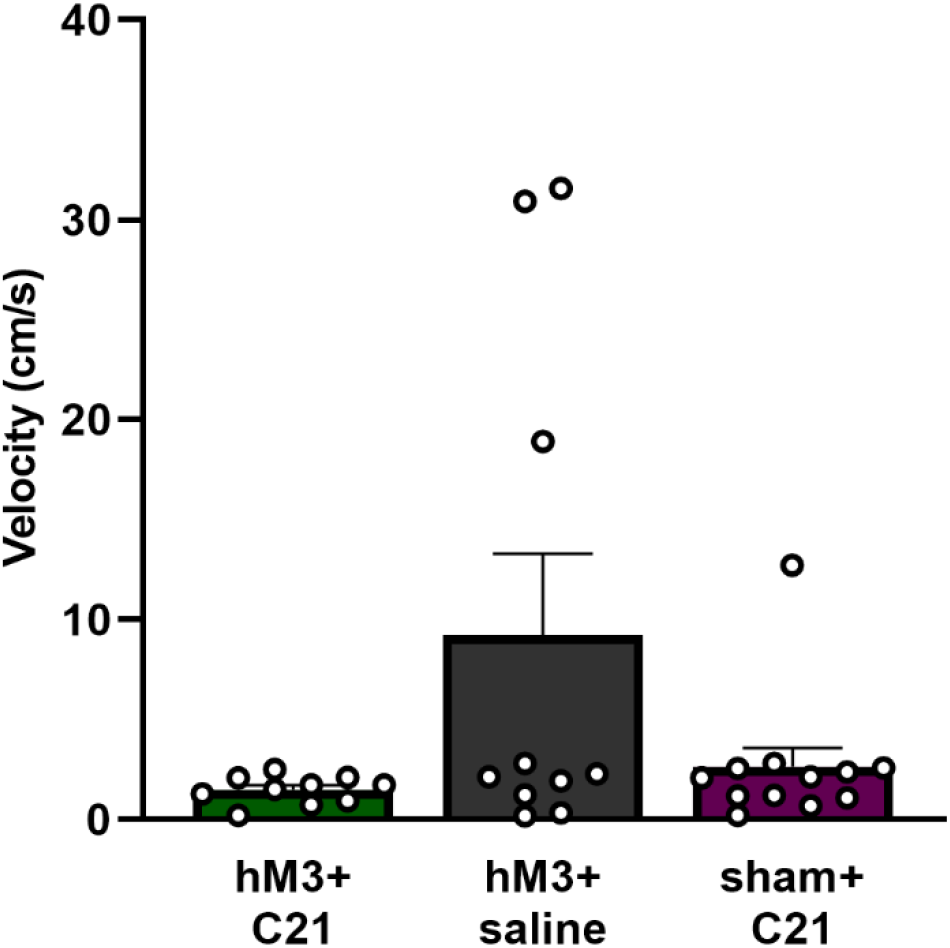
LS activation does not affect velocity. Velocity (centimeters/second; cm/s) of the hM3+C21 (green bars), hM3+saline (grey bars), and sham+C21 (purple bars) subjects during the social approach test. Data are presented as mean ± SEM. Dots represent individual data. (LMM, treatment effect: F_(2,29)_ = 3.21, *p* = 0.06).

**Figure S3.**
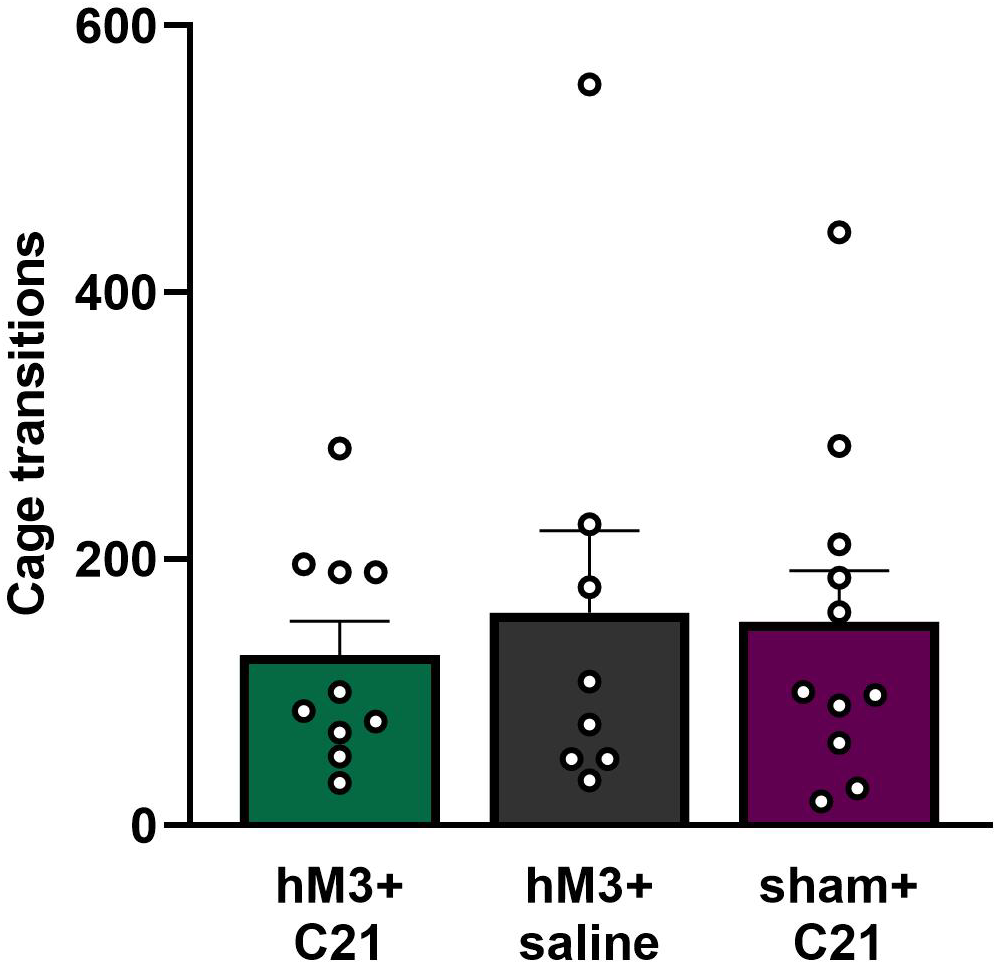
LS activation does not affect cage transitions. Cage transitions of the hM3+C21 (green bars), hM3+saline (grey bars), and sham+C21 (purple bars) subjects during the partner preference test. Data are presented as mean ± SEM. Dots represent individual data.

